# Telomere length, individual quality and fitness in female European starlings (*Sturnus vulgaris*) during breeding

**DOI:** 10.1101/416438

**Authors:** F. Criscuolo, M.F. Fowler, V.A. Fuhrer, S. Zahn, T.D. Williams

## Abstract

Telomeres, short guanine-rich repeats that cap linear chromosomes, are involved in cell senescence and organismal ageing. Our present understanding of telomere function oscillates between a marker of individual quality, which should be positively correlated with reproduction, to a marker of costs of reproduction (e.g. due to DNA damage). To test these ‘quality’ and ‘cost’ hypotheses, we tested the output of very simple predictions in a free-living population of European starlings *Sturnus vulgaris* where reproductive success of adult females was followed over 2 breeding seasons (with 2 broods per breeding). If telomere length indicate individual quality, low quality females (LQ, no fledgling) should have lower telomere lengths than high quality females (which produced fledglings in 1^st^ and 2^nd^ broods). Additionally, physiological determinants of adult individual quality (established in a previous study) and provisioning rate should be positively associated with adult telomere lengths. Finally, telomere length should predict future reproductive success. Adult telomere length was lower in LQ females but only during the chick-rearing period. Females producing larger clutches at fledging in the 1^st^ brood of the 1^st^ year also had longer telomeres. Provisioning rate was positively related to telomere length, as were plasma markers of oxidative damage, non-esterified fatty acids and triglycerides. Despite these associations, we found weak support for telomeres as indicators of individual quality. Telomere length failed in predicting future reproduction success and there was a lack of consistency in within-individual telomere length over the breeding season. In starlings, we suggest that telomere length may indicate current breeding capacities and energy status of female adults, rather than future fecundity/survival.

## Introduction

The fitness consequences of inter-individual variability in phenotype and genotype are a central concept in evolutionary biology (McNamara & Houston 1996; Wilson & Nussey 2010). This variability is important because evolutionary changes in a population depends on selection pressures that act on both phenotype and genotype. Therefore, inter-individual variability in fitness is a key parameter for selection and evolution to proceed, and its origin is one of the most fascinating questions in biology. In the literature, inter-individual variability has been described using many terms, such as “heterogeneity in fitness”, “frailty”, “health and vigor” or “organismal state”. More recently, it has commonly been described under the term of “individual quality” (Bergeron, Baeta, Pelletier *et al.* 2011). Individual quality is a much debated concept, which currently has no widely accepted formal definition (Wilson *et al.* 2010; Bergeron *et al.* 2011). However, there is some agreement on certain phenotypic or life-history traits that likely contribute to overall individual quality, *e.g.* timing of arrival at breeding grounds (Bauch, Becker & Verhulst 2013; Johnsen, Pauliny, Lifjeld *et al.* 2017), laying date or clutch size (Wilson *et al.* 2010), reproductive success (Bauch *et al.* 2013; Boonekamp, Dijkstra, Dijkstra *et al.* 2017), longevity and lifespan (Haussmann, Winkler & Vleck 2005; Pauliny, Wagner, Augustin *et al.* 2006; *Bize, Criscuolo, Metcalfe et al.* 2009; Barrett, Burke, Hammers *et al.* 2012; Boonekamp *et al.* 2017; Tricola, Simons, Atema *et al.* 2018).

In contrast, we still have a very poor understanding of physiological traits or mechanisms that might underpin variation in individual quality or life-history traits (Williams 2008; Wilson *et al.* 2010; Le Vaillant, Viblanc, Saraux *et al.* 2015). Telomeres are short guanine-rich repeated sequences that cap linear chromosomes and ensure their stability (Blackburn 2000). Those sequences progressively shorten with each cell division until they reach a minimum critical length, which will cause chromosome instability, cell senescence and cell death (Blackburn 2000). Until recently, because telomere loss mainly occurs during early life, studies on telomere dynamics have mainly focused on early developmental conditions, not adulthood (Geiger, Le Vaillant, Lebard *et al.* 2012; Monaghan & Spencer 2014; Nettle, Monaghan, Gillespie *et al.* 2015). However, several recent studies have suggested that telomeres might be more general biomarkers of adult individual quality (Hau, Haussmann, Greives *et al.* 2015; Le Vaillant *et al.* 2015; Bauch, Riechert, Verhulst *et al.* 2016) and might play a role in variation in life-history traits that contribute to fitness (Monaghan 2010; Monaghan *et al.* 2014; Ouyang, Lendvai, Moore *et al.* 2016; Johnsen *et al.* 2017). Measurement of telomere dynamics is therefore increasingly being used in ecological studies in order to investigate variation in life-history traits between individuals in wild populations (Monaghan & Haussmann 2006; Young 2018). Telomeres or telomere attrition rate have also been correlated to longevity among (Tricola *et al.* 2018; Wilbourn, Moatt, Froy *et al.* 2018) and within species, individuals with longer telomeres having a higher survival rate than those with shorter telomeres (Haussmann *et al.* 2005; Bize *et al.* 2009; Salomons, Mulder, van de Zande *et al.* 2009) but see (Caprioli, Romano, Romano *et al.* 2013). In relation to reproduction, telomere length has been correlated with the date of egg laying: adults of common terns (*Sterna hirundo*) with shorter telomeres arrived earlier in the breeding grounds and had higher reproductive performance (Bauch *et al.* 2013; Bauch, Becker & Verhulst 2014). Similarly, telomere length has been related to variation in foraging behaviour, individuals with shorter telomeres having higher foraging efficiency, which might mean that older individuals remain behaviourally flexible despite paying physiological costs (*Young, Kitaysky, Barger et al.* 2015). In king penguins (*Aptenodytes patagonicus*) and in common terns, a quadratic relationship between breeding success and telomere length has been reported: individuals with longer or shorter telomeres had higher breeding success than those with intermediate-length telomeres (*Bauch et al.* 2013; Le Vaillant *et al.* 2015). Both authors suggested that, (1) individuals with longer telomeres can increase their reproductive success paying less cost while individuals with short telomeres do poorly in reproduction and (2) reproduction remains costly in terms of telomere loss (Bauch *et al.* 2014), which in common terns could be related to provisioning rate of chicks (Bauch *et al.* 2013). Indeed, larger brood sizes were associated with greater rates of telomere loss in adults (Bauch *et al.* 2014; Reichert, Stier, Zahn *et al.* 2014). Finally, two independent studies reported assortative mating in relation to telomere length (Johnsen *et al.* 2017; Schull, Viblanc, Dobson *et al.* in press), suggesting that mate choice based on individual quality is somehow associated to telomere length. Therefore, our understanding of the role of telomere length in explaining individual fitness still oscillates from a marker of individual quality to a marker of DNA damage due to cost of life-history traits, such as reproduction. Most of these studies have simply correlated telomere length with life-history traits, and surprisingly, to date, very few studies have investigated this link in measuring both life-history and physiological traits (but see (Ouyang *et al.* 2016)) in order to better understand how telomere length and individual quality may be associated.

In this study, we investigated variation in telomere length in relation to individual quality in adult female European starlings (*Sturnus vulgaris*), integrating both life-history traits (*i.e.* current and future breeding success, survival) and physiological traits. We categorized females as low-, medium- and high-quality based on breeding productivity for between 1-4 broods over 1-2 years (see Materials and Methods). We predicted that, if relative telomere length is an inherent biomarker of “quality”, then, (1) high quality birds will have longer telomeres and low quality birds shorter telomeres when measured during incubation, (2) telomere length will be repeatable when measured between incubation and chick rearing for each individual, and (3) telomere length will be correlated with physiological measures of “quality” in starlings (*e.g.* high triglyceride, glucose, haematocrit, and haemoglobin levels, low oxidative stress, (Fowler & Williams 2017)), (4) we expect high quality individual (*i.e.* with longer telomeres) to successfully raise larger broods, sustaining a higher parental workload without suffering from higher physiological costs. Alternatively, if relative telomere length reflects costs of reproductive effort, then we predicted that (1) relative telomere length will decrease from incubation to brood 1 to brood 2, i.e. with increasing reproductive effort, (2) telomere length will be correlated with ultimate measures of parental workload, such as brood size at day 6-8, nest provisioning rate, as well as with (3) physiological markers of “workload” (e.g. oxidative stress).

## Material and Methods

### Species and area of study

We studied a nest-box breeding population of European starlings (*Sturnus vulgaris*) at Davistead Farm, Langley, British Columbia, Canada (49°10′N, 122°50′W) between April-June 2013 and females were followed in 2014 to measure survival and fecundity in the year after blood sampling. This site comprises around 150 nest boxes on farm buildings and mounted on posts around pastures. Nest boxes were checked daily from the first of April to determine laying date, clutch size, brood size at hatching, at day 6 post-hatching, and at fledging (day 21), and mass and tarsus of all chicks was recorded when chicks were 17 days old. We then collected the same data for second broods. All breeding females were captured at mid-incubation (between day 6 and 8) in their nest boxes by plugging the nest box hole before dawn, females were blood sampled (see below) and fitted with colour bands and individually numbered metal bands (Environment Canada permit # 10646; males were not captured or banded, and thus, identity of males is unknown).

During chick-rearing, females were caught using nest traps (Van Ert Enterprises, Leon, IA) as they entered nest boxes to feed chicks. All blood samples were collected by puncturing the brachial vein with a 26 ½-gauge needle and collecting blood (<700 μL) into heparinised capillary tubes. At the same time, fresh blood was collected for (1) hematocrit and hemoglobin measurements, (2) two blood smears were prepared for reticulocyte counts, and (3) glucose levels (mmol.L^-1^) were measured with a handheld glucose meter (Accu-chek Aviva®). Although females were captured using different methods, all birds were blood sampled within 3 minutes of being handled, and the only difference was that incubating birds were passively held in their nest box (mean 56 min., maximum 132 min. before being removed and blood sampled). However, there was no effect of time in box on plasma corticosterone levels in incubating females (r = 0.05, p > 0.73) suggesting that birds did not perceive this as a stressor. Blood samples were stored at 4°C for up to 4 h before being centrifuged for 6 min at 10,000 g, then plasma and red blood cells were separated and stored at −80°C until assayed. All research was conducted under Simon Fraser University Animal Care permits (# 657B-96,829B-96, 1018B-96).

### Determination of individuals’ quality

Individual quality of breeding females was assessed through a set of life history traits (including current and future breeding productivity, survival over 2 years) and was defined as following for mean brood size at fledging for each category and female sample size; see Results for statistical analysis):

1. Low quality females: had total brood failure (brood size at fledging = 0) for first broods, didn’t initiate a second brood in the same year and didn’t return the following year (n = 9, n = 5 and n = 0 during the incubation, chick-rearing and second brood phases, respectively);
2. Medium quality females: had average breeding productivity for first broods, attempted a second brood but failed (brood size at fledging for second brood = 0), and didn’t return the following year (n = 11, n = 9 and n = 0);
3. High quality females: had above-average breeding productivity for first broods, attempted a second brood (in which most chicks fledged), returned the following year and reared chicks from first and/or second brood (n = 6, n = 7 and n = 5).

### Measurement of physiological traits

As described above, plasma was extracted from blood samples in order to measure multiple physiological traits using standard methods reported in a previous study (see Fowler and Williams 2017 for details). Due to variation in the amount of plasma we obtained, for some individuals there were not enough plasma to run all assays, thus sample sizes differ for some physiological traits. These traits represented four “functional groups” of traits which were (1) aerobic/metabolic capacity, (2) oxidative stress and muscle damage, (3) intermediary metabolism and energy supply, and (4) immune function.

### Telomere analysis

DNA was extracted using the Blood Quick Pure kit (NucleoSpin, Macherey-Nagel) following the user’s manual, modified for bird blood. Briefly, the protocol consisted in (1) cellular digestion of red blood cells, (2) purification of DNA on a silica column and (3) elution of DNA in 50 μL of elution buffer. The modification of the protocol was to use 3 μL of red blood cells in 197 μL of PBS instead of 200 μL of whole blood. We then checked quality and concentration of DNA (μg/μL) with a NanoDrop 1000 (Thermo Scientific) spectrophotometer (absorbance ratio A260/280. 1.7; A260/230. 1.8).

We used real-time quantitative PCR technique (hereafter referred to as qPCR), developed by (Cawthon 2002) and adapted to avian telomeres by (Criscuolo, Bize, Nasir *et al.* 2009). This method amplified telomeric DNA and expressed its quantity as a T/S ratio, where T was the quantity of telomeric DNA and S the quantity of a gene used as a reference and amplified in a separate reaction. The reference gene classically used for birds was GAPDH, but was not well suited to European starlings probably due to a mutation (Nettle, Andrews, Reichert *et al.* 2016). In order to identify the most appropriate reference gene we conducted preliminary analyses considering a variety of candidate reference gene primer pairs available in our laboratory. The most consistent amplification profile and cleanest melting curve was obtained using Eurofins Genomics primers targeting the recombination activating 1 (RAG1) gene (accession number: XM_014873522), which we selected as our reference gene. The selection of our reference gene was based on comparison of a panel of several candidate genes for European Starlings including the commonly used one GAPDH. RAG1 showed completely stable qPCR results indicative of non-variable copy number, and is conserved among vertebrates (Venkatesh, Erdmann & Brenner 2001).

qPCR reactions were performed on separate 96-wells plates (n=6 for telomere sequence amplification, n=6 for the non-variable in copy gene number (RAG1). Each well contained a total volume of 10 μL, including 5 μL of SYBR green tampon, 10 ng of DNA, and 500 nmol/L of primers for telomere assay and 200 nmol/L for PYG assay. Reverse and forward primers’ sequences for both telomeric and PYG genes were, respectively: Tel1b: 5′ – CGGTTTGTTTGGGTTTGGGTTTGGGTTTGGGTTTGGGTT-3′; Tel2b: 5′ –GGCTTGCCTTACCCTTACCCTTACCCTTACCCTTACCCT-3′; RAG1-F: 5′ – TGCAAAGAGATTTCGATATGATG −3′; RAG1-R : 5′ – TCACTGACATCTCCCATTCC −3′. Telomeres thermal profile consisted of 2 min at 95°C (activation of the Taq polymerase), followed by 27 cycles of 15 seconds at 95°C (DNA denaturation), 30 seconds at 58°C (primers hybridization) and 30 seconds at 72°C (DNA amplification). RAG1 thermal profile was 2 min at 95°C, followed by 35 cycles of 15 sec at 95°C, 30 sec at 56°C and 1 min at 72°C. Both assays were followed by a melting curve analysis consisting in an increase of 1°C every 5 seconds (from 58°C to 95°C), to check for single peak amplification and the absence of primer-dimer artefact. Each sample was assayed in duplicate, and the mean of the two samples was used for analyses. Samples were randomly assigned over the 6 plates of telomere amplification in order to avoid a potential plate effect (an identical plate design was followed for the 6 PYG amplification plates). Each plate included a serial dilution (both telomeres and PYG) of 40 ng, 20 ng, 10 ng, 5 ng, 2.5 ng and 1.25 ng DNA. This dilution was made with a pool of DNA from the most concentrated samples. This dilution was needed to generate a reference curve on each plate to control for the amplifying efficiency of the qPCR, which ranged from 94 to 109 % for PYG and from 99 to 104 % for telomeres (an efficiency of 100 % corresponded to a doubling of DNA at each cycle, the closer to 100 % is the better). Each plate also contained 4 different samples chosen randomly which were repeated over all runs to check for repeatability in amplification values. Intra-plate and inter-plate coefficients of variation (based on Cq values) were 0.34 ± 0.03 % and 0.64 ± 0.07 % for the PYG gene assay and 1.92 ± 0.22 % and 2.10 ± 0.71 % for the telomere assay, respectively. Intra-plate and inter-plate coefficients of variation (based on T/S ratio) were 10.64 ± 0.84 % and 11.77 ±4.59 %, respectively.

We used equation 1 from (Pfaffl 2001), which was a mathematical model of relative expression ratio in real-time quantitative PCR allowing the utilisation of each plate efficiency (and not a mean value over all runs) to calculate the relative telomere length. The ratio of the telomere gene was expressed in a focal sample versus a control sample in comparison to the reference gene RAG1. We used the following equation: ratio T/S = [(E_telomere_)ΔCq_telomere_(Control-sample)]/[(E_PYG_)ΔCq_PYG_(Control-Sample), where E corresponded to the efficiency of the reaction, Cq_telomere_ corresponded to the number of amplification cycles recorded to reach the crossing point with the fluorescence measurement threshold for the telomere sequence, Cq_PYG_ for the control gene. ΔCq corresponded to the difference between the Cq of the control sample and the focal sample.

Relative telomere length is then calculated as a ratio between the number of cycles of amplifications needed to reach the detection threshold for the RAG1 gene (Cq RAG1) and those needed for the telomere sequence amplification (Cq Tel), all corrected for the actual efficiencies of the amplification reactions. As ratio utilisation in statistics may not always follow a normal distribution and often necessitates log-transformation, we tested the outcome of using instead the residuals of the linear regression between Cq RAG1 and Cq Tel obtained for each individuals. The rationale is that, for instance, high residuals will correspond to a higher number of amplification cycles of the telomere sequence needed to reach the detection threshold (and then a shorter telomere length), for a given quantity of DNA (Cq RAG1). All analyses has been duplicated to test whether the final outcome is illustrating the same significant relationship between physiological and fitness parameters and telomere length, as calculated using the log-transformed T/S ratio (see Electronic Supplementary Material, ESM1).

### Statistical analysis

Data were analysed using R version 3.3.2 (R Core Team 2016) or SPSS 18.0. We tested for normality with both a Shapiro-Wilk’s test and QQ plots distribution, and when data were not normal, we used log10 transformation (e.g. T/S ratio). Since we studied inter-individual variability, we didn’t remove any data as statistical outliers. All effects with P < 0.05 were considered as significant. Post-hoc comparisons were done using Bonferroni correction. Differences in brood size at fledging in the first year, and in total fledging number over the two years among females of different qualities were tested using a Multivariate General Linear Model (MANOVA) with Individual Quality as fixed factors.

Our statistical analysis was based on the following steps: (i) we tested how telomere length is related to individual quality using a Mixed Model with log-transformed T/S ratio as the response variable, and female Individual Quality and Breeding Stages as fixed factors. To control for repeated measurements, female Identity was included as random factor; (ii) to test relationships between physiological variables and individual telomere length we used Principal Component analysis on seven physiological variables previously identified as measures of physiological state and costs of reproduction (Table 2A; see (Fowler *et al.* 2017) for more details). We used the individual PCA1, PCA 2 and PCA 3 values as predictors of log-transformed T/S ratio using a General Linear Model (GLM). Only individual physiological and telomere values recorded during the chick rearing phase were used in this step; (iii) finally, sequential GLM were run to test how log-transformed T/S ratio predicts current reproductive effort (provisioning rate and number of chicks fledged from the 1^st^ brood), future reproduction (initiating a second brood in year 1, return rate to year 2, number of chicks fledged in subsequent breeding attempts) and total reproductive success (total number of fledgings over 2 years). Depending on the response variable, appropriate covariates were included to control for among-individual variability in reproduction success (see Results).

To test whether relative telomere length could be calculated following residuals of the correlation between Cq RAG1 and Cq Tel, we duplicated the statistical analysis using the residuals of the linear regression between Cq RAG1 and Cq Tel. This is described in ESM1.

## Results

### Individual quality and breeding productivity

Individual quality was based on breeding success, measured through brood size at hatching during the first year, and at fledging over two years and up to four broods. All three variables were significantly different among LQ, MQ and HQ females (MANOVA, Roy’s largest root, D_3, 22_ = 45,221, P < 0.001), *e.g.* mean cumulative number of chicks fledged over 2 years was zero, 4.8 ± 0.6, and 12.6 ± 1.1, respectively. Bonferroni post-hoc comparisons showed that LQ females have lower brood size at the three stages when compared to the two other groups (model estimates: 1^st^ year brood size at incubation: −4,94 ± 1.24, P = 0.001); 1^st^ year fledging: −4,00 ± 0.87, P < 0.001; total fledging number over 2 years: −2,65 ± 0.74, P = 0.001). HQ females had higher fledging brood sizes than MQ during the 1^st^ year (2,80 ± 0.93, P = 0.006) and in total over the 2 years of the study (7,66 ± 1.32, P < 0.001), but not at 1^st^ year’ incubation stage (0,59 ± 0.78, P = 0.46).

### Telomere length and individual quality

Figure 1A illustrates within-individual variation in telomere length over the entire study period, showing a large individual variability. Table 1 shows the results of the GLMM testing for difference in relative telomere length in relation to variation in individual quality. There was no significant main effect of breeding stage on relative telomere length, although LQ females were characterized by a close-to-significant decrease in telomere length between incubation and chick-rearing (Figure 1B, Table 1). However, there was a significant individual quality x breeding stage interaction: relative telomere length varied with individual quality, but only when measured during chick-rearing. LQ females had shorter telomeres than MQ and HQ females at the chick rearing stage, while no differences were significant at the preceding incubation stage (Figure 2, Table 1).

**Table 1.** Results of a Mixed Model with Log10-transformed relative telomere length (TS ratio) as a response variable, with individual quality and breeding stages as fixed factors and female identity as random factor. HQ = High Quality, MQ = Medium Quality, LQ = Low Quality. Post-hoc comparisons were done using Bonferroni corrections, among groups during the incubation and chick rearing stages, and within groups between incubation and chick rearing stages. Significant P values are indicated in bold. Residuals of each models followed a normal distribution (checked using Kolmogorov-Smirnov test and QQ plot).

| Response variable: **Log10 (TS ratio)** | Estimates | D.F | F | P |
| --- | --- | --- | --- | --- |
| Intercept | -0.153 ± 0.048 | 1, 25 | 58.336 | **<0.001** |
| **Individual Quality** |  | 2, 25 | 10.158 | **0.006** |
| MQ females vs. HQ females | 0.135 ± 0.091 |  |  | 0.411 |
| MQ females vs. LQ females | 0.250 ± 0.083 |  |  | **0.008** |
| HQ females vs. LQ females | 0.116 ± 0.114 |  |  | 0.933 |
| **Breeding stage (Chick rearing vs. Incubation)** | -0.042 ± 0.071 | 1, 25 | 1.887 | 0.170 |
| **Individual Quality x Breeding stage** | 1.524 ± 0.317 | 2, 25 | 7.170 | **0.028** |
| During Incubation |  |  |  |  |
| MQ females vs. HQ females | 0.227 ± 0.163 |  |  | 0.931 |
| MQ females vs. LQ females | 0.064 ± 0.111 |  |  | 1.000 |
| HQ females vs. LQ females | -0.164 ± 0.185 |  |  | 0.999 |
| During Chick rearing |  |  |  |  |
| MQ females vs. HQ females | 0.042 ± 0.066 |  |  | 1.000 |
| MQ females vs. LQ females | 0.437 ± 0.126 |  |  | **0.008** |
| HQ females vs. LQ females | 0.395 ± 0.124 |  |  | **0.021** |
| Between Incubation & Chick rearing |  |  |  |  |
| Among HQ females | 0.143 ± 0.154 |  |  | 0.999 |
| Among MQ females | -0.042 ± 0.071 |  |  | 1.000 |
| Among LQ females | -0.416 ± 0.153 |  |  | 0.079 |

**Figure 1:**
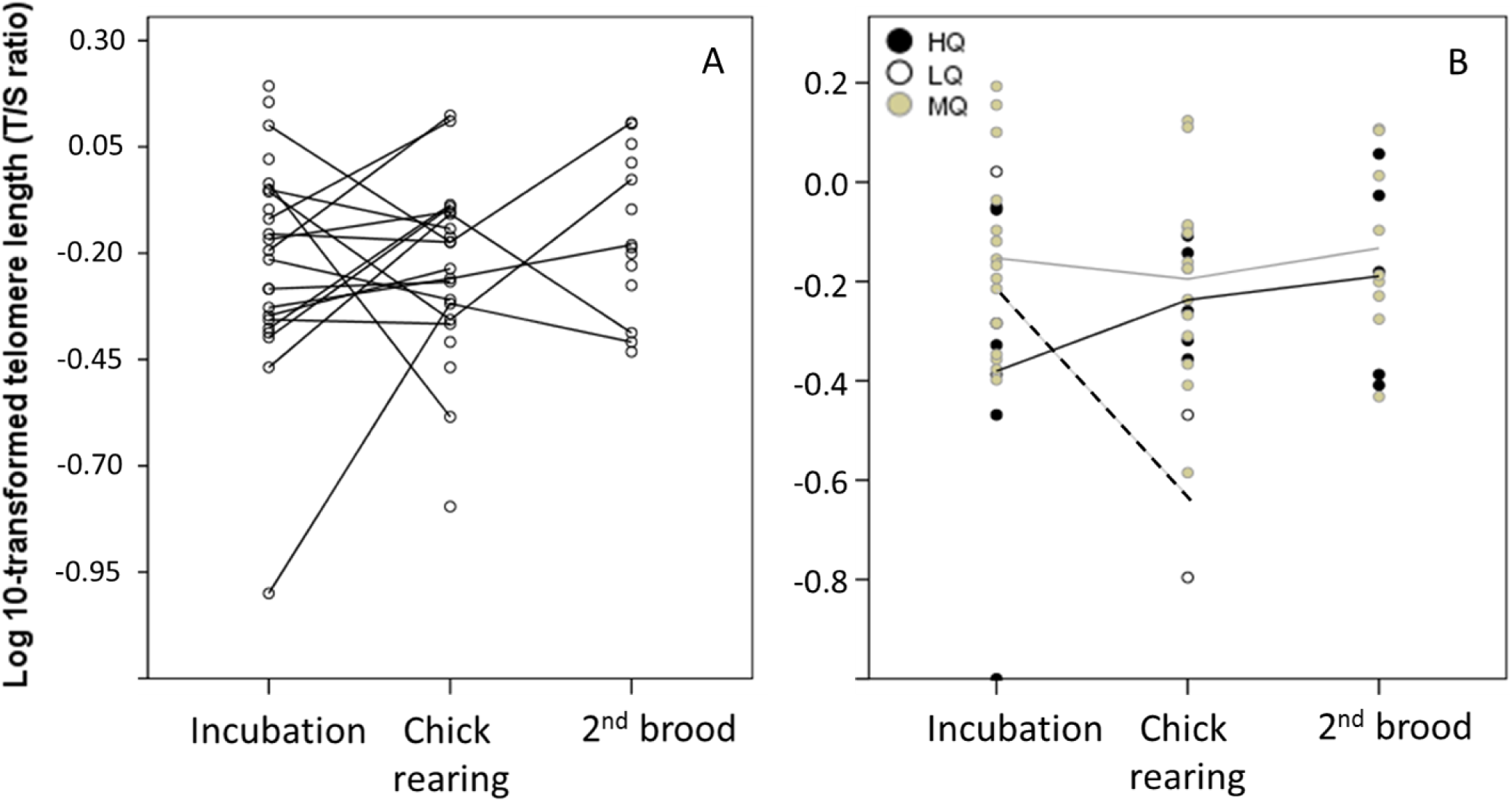

**Figure 2:**
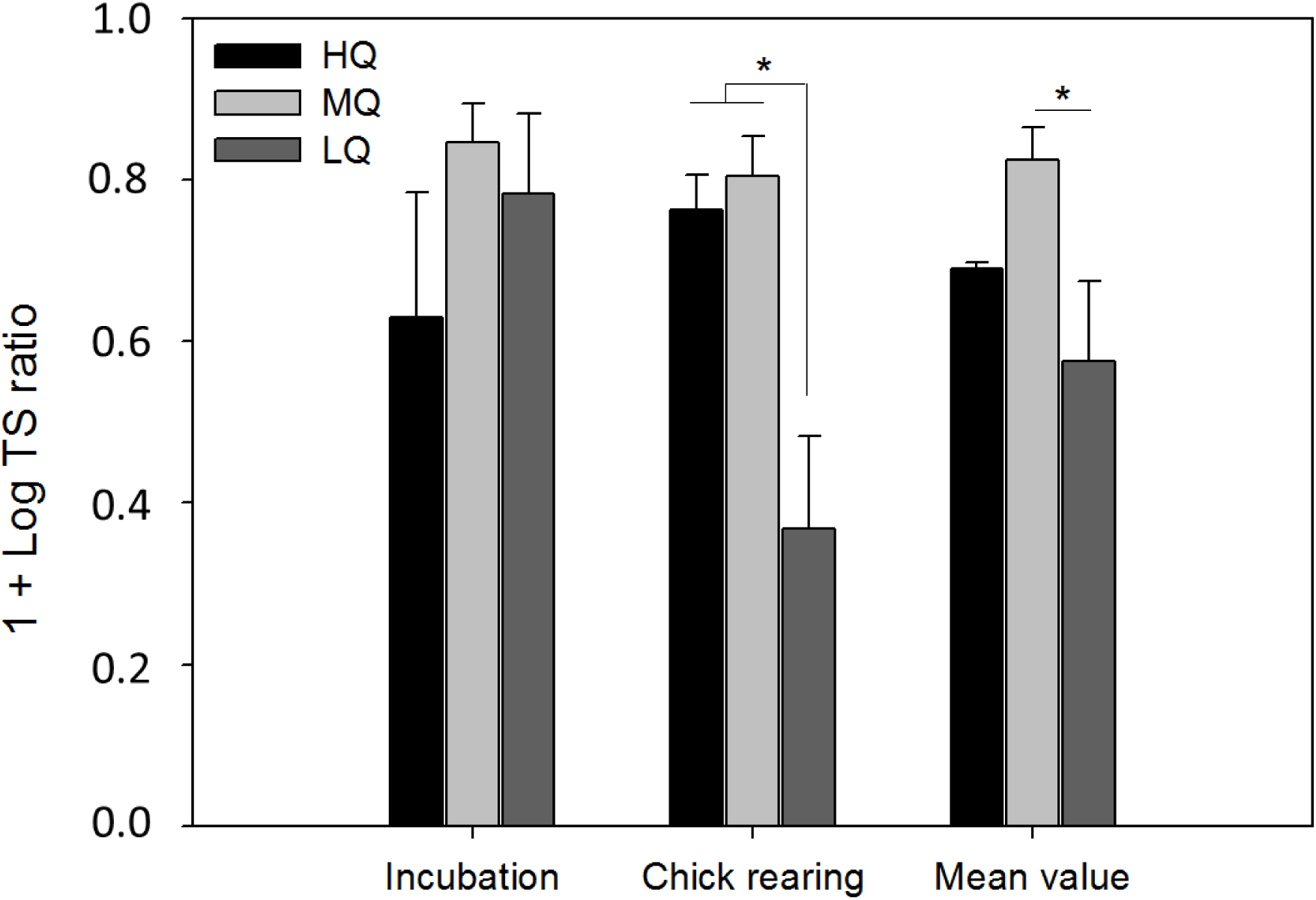
Differences in relative telomere length over reproduction stages of starling adult females in relation to their individual quality (HQ: High quality, N = 10; MQ: Medium quality, N = 29; LQ: Low quality, N = 5). Relative telomere length is expressed as 1 + log10-transformed T/S ratios obtained from qPCR amplification (see Methods). Telomeres were significantly shorter in LQ females during chick rearing than in HQ and MQ females. Overall mean values of telomere length differed significantly between Medium and Low Quality females (see Table 2). * P < 0.05. See Table 2 for statistics.

**Table 2.** A. Variables forming the axes (PCA1, PCA2 and PCA3) of a Principal Component Analysis conducted on 7 physiological traits measured in adult females during the chick-rearing breeding stage, and previously shown to be related to cost of reproduction in starlings (Fowler and Williams 2017). Only eigenvalues over 0.4 were taken into account. Total variance explained by the model: 73.8%. B. General Linear Model testing for relationships between Log10-transformed relative telomere length (TS ratio) and the individual PCA values obtained from 7 physiological traits measured in adult females during the chick-rearing breeding stage. Residuals followed a normal distribution (checked using Kolmogorov-Smirnov test and QQ plot). Significant results are given in bold.

| **A** | PCA1 | PCA2 | PCA 3 |
| --- | --- | --- | --- |
| **Physiological variables** |  |  |  |
| Hematocrit | 0.850 |  |  |
| Haemoglobin | 0.834 |  |  |
| Uric acid | 0.777 |  |  |
| Triglyceride |  | 0.806 |  |
| dROMs |  | 0.768 |  |
| NEFA |  | 0.608 |  |
| Haptoglobin |  |  | 0.943 |
| **% variance explained** | **35.6** | **21.0** | **17.8** |

| **B** | Estimates | D.F | F | P |
| --- | --- | --- | --- | --- |
| Response variable: **Log10 (TS ratio)** |  |  |  |  |
| Intercept | -0.265 ± 0.047 | 1, 17 | 31.908 | **<0.001** |
| PCA 1 | -0.035 ± 0.047 | 1, 17 | 0.538 | 0.475 |
| PCA 2 | 0.129 ± 0.048 | 1, 17 | 7.133 | **0.018** |
| PCA 3 | -0.020 ± 0.048 | 1, 17 | 0.174 | 0.683 |

### Telomere length and physiological traits

Principal Component Analysis conducted using the 7 physiological traits measured during chick-rearing enables us to discriminate three physiological axes (PCA1, PCA2 and PCA3) explaining 74% of the total variance (Table 2A). Only PCA2 was significantly correlated to relative telomere length (Table 2B), indicating that females with longer telomeres had higher plasma levels of triglycerides, NEFA and dROMs (Figure 3).

**Figure 3:**
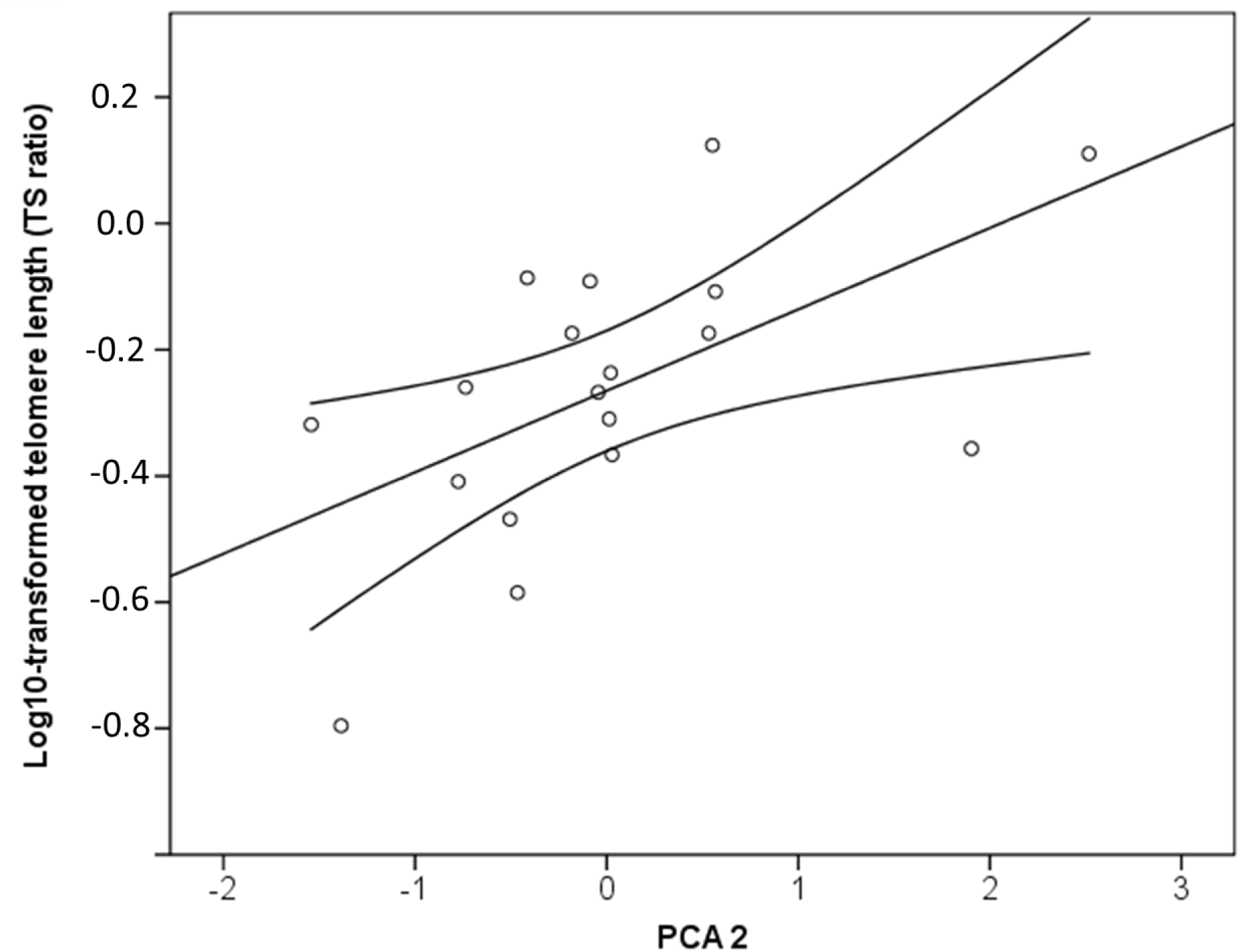
Pearson’s linear regression checking for linear relationship between individual telomere length of adult females (TS ratio log10 transformed) and their PCA 2 values, all measured during the chick rearing breeding stage (see Methods for details). Individuals with high PCA2 values have higher plasma levels of non-esterified fatty-acids, triglycerides and dROMs (Table 2A). Lines represent 95% confidence interval. Log10 (TS ratio) = 0.129 (PCA2) – 0.265, r^2^ = 0.37, P = 0.013. See Table 2 for detailed statistics.

### Telomere length and breeding effort in the first year

Relative telomere length was positively related to brood size at fledging: females with longer telomeres successfully raised a larger number of chicks (Figure 4, Table 3A). During chick-rearing, provisioning rate was marginally positively correlated to telomere length (Table 3A). Individuals with longer telomeres have a higher nest provisioning rate, and conversely, individuals with short telomeres have lower provisioning rates (Figure 5).

**Table 3.** Results of General Linear Models (with a logistic distribution for return rate and 2^nd^ brood initiation) testing for relationships between Log10-transfromed telomere length (TS ratio) of adult starling females and: (A) current effort of reproduction, (B) future prospects of reproduction during the same breeding season (2013, 2^nd^ brood), (C) future prospects of reproduction during the next breeding season (2014), (D) total reproduction success over 2013 and 2014. In B and C models, first brood size (in 2013 and 2014) and first year total fledging number (in 2013) were included as a covariates to control for the initial brood or first year breeding costs. Adult telomere length was measured during the chick rearing period of the first brood. Significant results are indicated in bold.

|  |  |  |  |  |
| --- | --- | --- | --- | --- |
| **A. Current breeding effort (2013)** |  |  |  |  |
| Response variable: *Brood size at day 17* | Estimates | D.F | F | P |
| Intercept | 4.010 ± 0.520 | 1, 17 | 7.707 | **<0.001** |
| Log10 (TS ratio) | 3.510 ± 1.613 | 1, 17 | 2.176 | **0.044** |
| Response variable: *chick provisioning rate* | Estimates | D.F | F | P |
| Intercept | 5.667 ± 0.765 | 1, 17 | 7.409 | **<0.001** |
| Log10 (TS ratio) | 4.953 ± 2.371 | 1, 17 | 2.088 | **0.052** |
| 1<sup>st</sup> brood size | 0.170 ± 0.365 | 1, 17 | 0.465 | 0.648 |
| **B. Same year breeding prospects (2013)** |  |  |  |  |
| Response variable: *Initiation of a 2<sup>nd</sup> brood* | Estimates | D.F | F | P |
| Intercept | 0.866 ± 1.553 | 1, 17 | 0.558 | 0.577 |
| Log10 (TS ratio) | 2.007 ± 2.608 | 1, 17 | 0.769 | 0.442 |
| 1<sup>st</sup> brood size | -0.016 ± 0.336 | 1, 17 | -0.048 | 0.962 |
| Response variable: *2<sup>nd</sup> brood size at fledging* | Estimates | D.F | F | P |
| Intercept | -0.025 ± 1.123 | 1, 17 | -0.022 | 0.983 |
| Log10 (TS ratio) | -1.031 ± 1.858 | 1, 17 | -0.555 | 0.587 |
| 1<sup>st</sup> brood size | 0.145 ± 0.247 | 1, 17 | -0.588 | 0.565 |
| **c. Next year breeding prospects (2014)** |  |  |  |  |
| Response variable: *adult female return rate* | Estimates | D.F | F | P |
| Intercept | -3.381 ± 1.667 | 1, 17 | 1.667 | **0.043** |
| Log10 (TS ratio) | -1.076 ± 3.034 | 1, 17 | -0.354 | 0.723 |
| 1<sup>st</sup> year fledging number | 0.494 ± 0.300 | 1, 17 | 1.650 | 0.099 |
| Response variable: *2<sup>nd</sup> year, 1<sup>st</sup> brood size at fledging* | Estimates | D.F | F | P |
| Intercept | -0.755 ± 1.297 | 1, 17 | -0.582 | 0.569 |
| Log10 (TS ratio) | -1.406 ± 2.430 | 1, 17 | -0.555 | 0.570 |
| 1<sup>st</sup> year fledging number | 0.450 ± 0.232 | 1, 17 | 1.944 | 0.070 |
| Response variable: **2<sup>nd</sup> year, initiation of a 2<sup>nd</sup> brood** |  |  |  |  |
|  | Estimates | D.F | F | P |
| Intercept | -0.025 ± 1.123 | 1, 17 | -0.022 | 0.983 |
| Log10 (TS ratio) | -1.031 ± 1.858 | 1, 17 | -0.555 | 0.587 |
| 1<sup>st</sup> year fledging number | 0.145 ± 0.247 | 1, 17 | -0.588 | 0.565 |
| Response variable: **2<sup>nd</sup> year, 2<sup>nd</sup> brood size at fledging** |  |  |  |  |
|  | Estimates | D.F | F | P |
| Intercept | 0.649 ± 0.403 | 1, 17 | 1.610 | 0.128 |
| Log10 (TS ratio) | 1.008 ± 0.754 | 1, 17 | 1.336 | 0.201 |
| 1<sup>st</sup> year fledging number | -0.124 ± 0.079 | 1, 17 | -1.557 | 0.140 |
| 2<sup>nd</sup> year, 1<sup>st</sup> brood fledging number | 0.253 ± 0.077 | 1,17 | 3.290 | **0.005** |
| Response variable: **2<sup>nd</sup> year total fledging number** |  |  |  |  |
|  | Estimates | D.F | F | P |
| Intercept | -0.298 ± 1.670 | 1, 17 | -0.178 | 0.861 |
| Log10 (TS ratio) | -0.754 ± 3.129 | 1, 17 | -0.241 | 0.813 |
| 1<sup>st</sup> year fledging number | 0.441 ± 0.298 | 1, 17 | 1.479 | 0.159 |
| **D. Total breeding success (2013 and 2014)** |  |  |  |  |
| Response variable: **Total number of fledging over 2 years** |  |  |  |  |
|  | Estimates | D.F | F | P |
| Intercept | 6.283 ± 1.472 | 1, 17 | 4.269 | **<0.001** |
| Log10 (TS ratio) | 3.553 ± 4.564 | 1, 17 | 0.778 | 0.447 |

**Figure 4:**
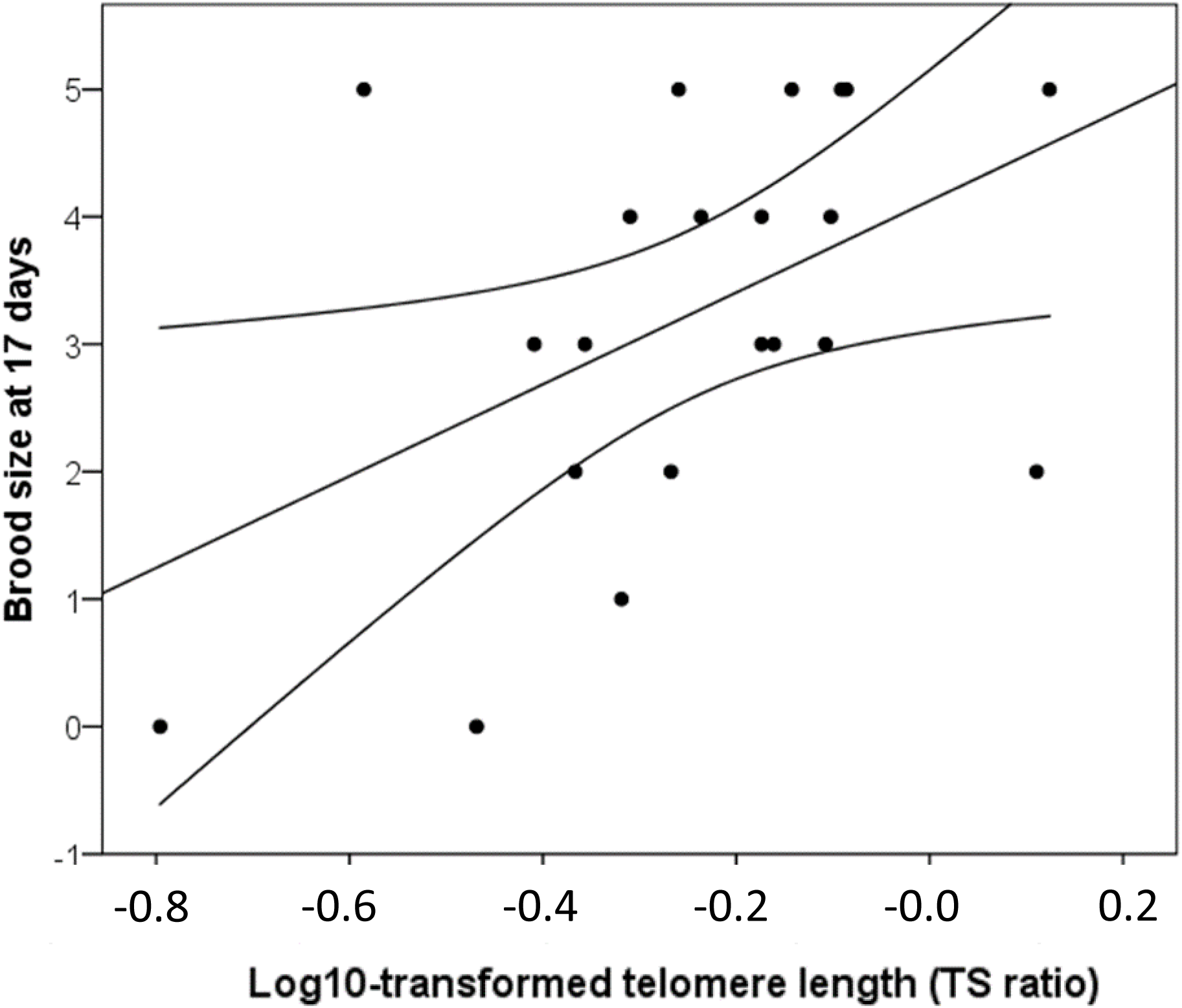
Pearson’s linear regression checking for linear relationship between individual telomere length of adult females (TS ratio log10 transformed) and their 1^st^ brood size in 2013. TS ratio was measured during the chick rearing breeding stage (see Methods for details). Lines represent 95% confidence interval. Brood size = 3.601 (Log10 (TS ratio)) + 4.126, r^2^ = 0.29, P = 0.029. See Table 3 for detailed statistics.

**Figure 5:**
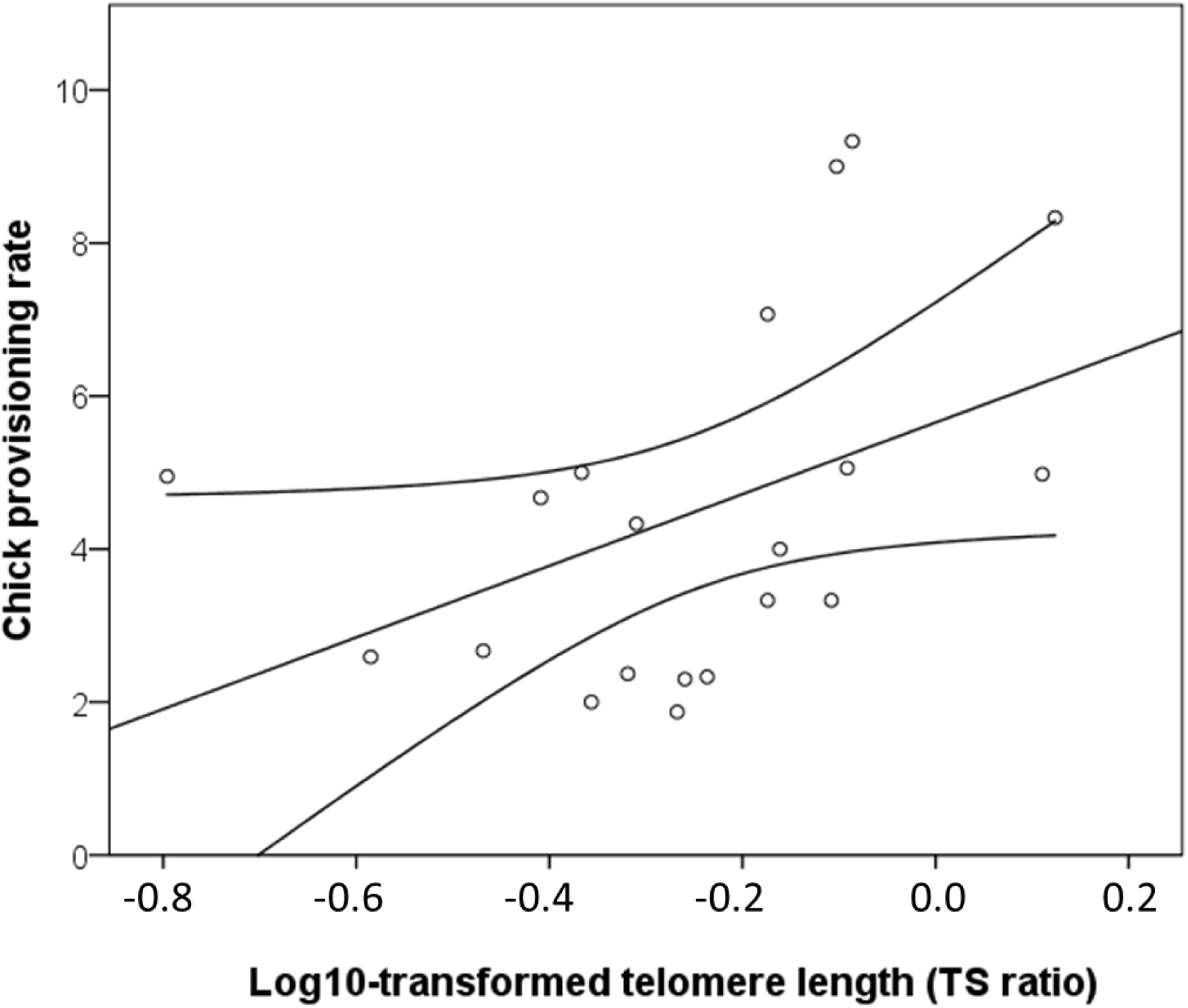
Pearson’s linear regression checking for linear relationship between individual telomere length of adult females (TS ratio log10 transformed) and the females’ provisioning rate of their 1^st^ brood size in 2013. TS ratio was measured during the chick rearing breeding stage (see Methods for details). Lines represent 95% confidence interval. Provisioning = 4.682 (Log10 (TS ratio)) + 5.654, r^2^ = 0.19, P = 0.054. See Table 3 for detailed statistics.

### Telomere length, breeding prospects and total breeding success

Relative telomere length of females at chick-rearing during 1^st^ broods did not predict the probability of females initiating a 2^nd^ brood during the first year of breeding or brood size at fledging of this 2^nd^ breeding attempt (Table 3B). Similarly, when looking at the following breeding season (2^nd^ year), return rate, 1^st^ brood size at fledging, initiation of a 2^nd^ brood or 2^nd^ brood size at fledging were all independent of relative telomere length in the preceding year (Table 3C). Finally, relative telomere length was not a proxy of total breeding success in terms of number of chicks fledged over the two years of the study (Table 3D).

## Discussion

We investigated variability in relative telomere length in female adult European starlings during three different stages of reproduction: incubation, chick-rearing, and second brooding. We found some support for our first main hypothesis: telomere length was related to individual quality as determined by overall breeding productivity, HQ and MQ females had longer telomeres than LQ females during the chick-rearing stage. However, we had weak support for our second hypothesis, finding a low within-individual repeatability over breeding stages (and only for MQ females), while individual variation in relative telomere length independently of individual quality and breeding stage was substantial. However, LQ females were close to have a significant shortening of their telomeres between incubation and chick-rearing stages. We found mixed support for our third prediction since, among the 7 physiological traits we comprehensively measured to characterize individual physiological state (including aerobic capacity, immune function, oxidative stress, and energy metabolism), only three of the physiological traits measured were linked to telomere length during chick-rearing of the first brood: NEFAs, triglycerides and reactive oxygen metabolites concentrations (dROMs) were positively correlated with relative telomere length. Finally, relative telomere length predicted individual variation in brood size at fledging, but only in the current breeding attempt. Chick provisioning rate was positively related to female telomere length, supporting the idea that females with longer telomeres might be able to support higher rates of workload. However, we found little support for the hypothesis that telomere length was related to individual quality based on future fecundity or survival (*i.e.* ultimate measures of cost of reproduction).

We predicted that HQ individuals would have longer telomeres and, conversely, LQ individuals would have shorter telomeres. This view is based on the simple idea that telomere length reflects the lifestyle of an individual, characterizing the cumulative impact of stress, or more comprehensively the higher sensitivity to stress of LQ individuals (Monaghan *et al.* 2006). However, previous studies did not always corroborate this idea, rather finding a curvilinear relationship between reproduction success and telomere length (Bauch *et al.* 2013). In our study, telomere length was significantly lower only in LQ females, which had zero reproductive success. However, no marked difference in telomere length was found between HQ and MQ females, even though there was marked variation in cumulative breeding productivity in our sample of LQ, MQ and HQ adult females (zero, 4 and 12 fledged chicks respectively over 1–2 years). In our study, individuals with longer telomeres had higher reactive oxygen metabolites concentrations. That could mean that reactive oxygen metabolites are not linked to telomere attrition which would contradict a recent review of the literature (Reichert & Stier 2017). However, an interesting recent theoretical view about the trade-offs leading to telomere shortening is that telomere maintenance is costly *per se* (due to energy cost of cellular and DNA processing, (Young 2018)), opening the possibility that, for instance, the LQ females are unable to pay the cost of telomere maintenance independently of the oxidative balance issue whereas the MQ and HQ females do. Secondly, our results could suggest that there is no link between parental workload and oxidative stress levels or telomeres in our birds which seems to be unlikely (see below). Thirdly, individuals with long telomeres are either able to better cope with reactive oxygen metabolites because they pay less shortening cost, or because they are able to restore any telomere loss. Nevertheless, we acknowledge a methodological limitation of qPCR, which does not allow us to distinguish the proportion of long telomeres of individuals. Reproduction costs may affect in priority long telomeres, and that LQ females suffered from a reduction of their proportion of long telomeres which has been previously suggested to better explain individual variation in fitness (Bauch *et al.* 2014). It would be interesting in the future to test how oxidative stress levels correlates with long telomeres’ shortening rate.

We found a positive relationship between females’ telomere length and parental effort in terms of brood size and nest provisioning rate during chick-rearing. Increasing reproductive effort has been suggested to lead to an increase in metabolic rate (Barnes & Partridge 2003) and oxidative stress (Stier, Reichert, Massemin *et al.* 2012; Colominas-Ciuró, Santos, Coria *et al.* 2017; Merkling, Blanchard, Chastel *et al.* 2017), particularly in other bird species, which should be associated with shorter telomeres. However, the positive relationship between telomere length and provisioning rate in our study is consistent with Young et al’s (2015) suggestion that telomere length has widespread predictive value for determining foraging behavior of individuals. They found that telomere length was related to foraging depth, maximum depth and foraging efficiency in thick-billed murres (*Uria lomvia*), although follow-up study in the same species by Young et al. (2016) found no relationship between telomere length, colony attendance and foraging trip rate. A positive relationship between telomere length and parental effort is more consistent with life-history theory; individuals should not work hard enough to kill themselves and will trade-off self-maintenance and reproductive investment. Higher-quality birds with longer telomeres might be able to increase their reproductive effort in order to increase their fitness without paying costs, either because they are more experienced (i.e. age-dependent relationship between reproductive investment and metabolic costs, (Speakman & Garratt 2014)) or because they buffer themselves and their chicks against deleterious effects of high metabolism (i.e. shielding hypothesis, (Blount et al. 2015)). On the contrary, LQ females were the only ones showing a reduction of their telomere lengths between incubation and chick-rearing stages (p=0.079). This reduction is likely to be the main factor defining our quality group difference, and might be related to cost of (attempted) reproduction. LQ females may have been unable to maintain the cost of parental workload while feeding chicks such as oxidative stress associated with sustained high metabolic rate (Barnes *et al.* 2003). Numerous studies have shown that high metabolic rate is correlated with higher oxidative stress levels and therefore potentially also to telomere shortening (von Zglinicki 2002; Monaghan, Metcalfe & Torres 2009). Experimental studies, where parental effort and costs of reproduction are manipulated, are required to further test these ideas (Metcalfe & Monaghan 2013). Nevertheless, our data suggest that having long telomeres is associated with higher fledging success in the first breeding year, which remains a strong argument underlining that individual quality partly has a molecular and physiological bases (Le Vaillant *et al.* 2015). How might our result be reconciled with previous results showing that individuals with shorter telomeres actually performed better (Bauch *et al.* 2013; Bauch *et al.* 2014)? As suggested by Bauch et al., telomeres may reflect past reproductive costs (*i.e.* parental effort over several breeding seasons) in common terns, which may be stronger in driving telomere links with fitness in a bird living for decades, while in a short-lived species as the European starling, telomere length measured when reproduction starts is less likely to compute cumulated past reproductive costs.

An alternative explanation for individuals providing higher parental effort and suffering from higher oxidative damage but also having long telomeres is that they have specific DNA restoration abilities. For instance, the major known telomere restoration mechanism involves the enzyme telomerase (Blackburn 2009). Increased telomerase activity, for instance in response to stress, may have beneficial impact both on telomeres but also on individual physiological traits (Bernardes de Jesus & Blasco 2012; Criscuolo, Smith, Zahn *et al.* 2018). We didn’t measure telomerase activity in the current study but this would be worthwhile in future studies, given that we found relative telomere length increasing in some adult females between breeding stages (Figure 1A). While there is no evidence that starlings belong to a family of birds characterised by particular rate of change in telomere length over age (*i.e*. having specific telomere restoration abilities (Tricola *et al.* 2018)), we still lack accurate data focusing on potential acute changes in telomerase activity in response to stress (*Beery, Lin, Biddle et al.* 2012), as well as on individual variability in such a compensatory mechanism for the reproduction / self-maintenance trade-off.

Compensation mechanisms associated with relative telomere length could occur through changes in physiological traits linked to individual quality. In starlings, the physiological basis of costs of reproduction was previously identified with a suite of seven traits changing in response to handicapping (wing-clipping) as an experimental manipulation of parental effort. Females with lower future fecundity and lower return rates had lower oxygen-carrying capacity (lower hematocrit and hemoglobin levels), lower energy reserves (plasma non-esterified fatty acid and triglyceride levels), decreased immune function (lower haptoglobin levels), elevated levels of oxidative stress (higher levels of dROMs [reactive oxygen metabolites] and lower levels of the endogenous antioxidant uric acid (Fowler & Williams 2018). The fact that females with longer telomeres also have higher levels of two of these physiological markers of either feeding status or energy fuel mobilization (triglycerides, NEFA) suggests that those females were in better body condition than those with shorter telomeres. Since those measurements were done during the more demanding period of breeding (*i.e.* chick-rearing) and that females with longer telomeres engaged in higher parental effort and reared more fledglings, we conclude that telomere length reflects higher metabolic capacities rather than a reallocation of energy towards self-maintenance. One previous study highlighted that males common terns, which provide the largest part of chick provisioning, suffer higher telomere shortening and had higher corticosterone plasma levels, while no link was found in females (Bauch *et al.* 2016). In accordance, we found no link between telomere length and corticosterone levels in our females (data not shown), but we do know that females do most of chick provisioning in starlings (Fowler and Williams 2015). Whether females are better protected from adverse effects of corticosterone on telomeres (e.g. due to estrogens (Tissier, Williams & Criscuolo 2014)) remains to be tested.

Several recent studies have suggested that telomeres might be general biomarkers of individual quality in adults (Hau *et al.* 2015; Le Vaillant *et al.* 2015; Bauch *et al.* 2016) and might play a role in variation in life-history traits that contribute to fitness (Monaghan 2010; Monaghan *et al.* 2014; Ouyang *et al.* 2016; Johnsen *et al.* 2017). For example, in king penguins *Aptenodytes patagonicus*, birds with longer telomeres arrived earlier for breeding at the colony, and had higher breeding performances than individuals with shorter telomeres (Le Vaillant *et al.* 2015). In contrast, in tree swallows *Tachycineta bicolor,* males and females with longer telomeres had lighter nestlings (Ouyang et al. 2016). In our study, we had comprehensive fitness-related measures in terms of current breeding productivity, future fecundity and survival, with total cumulative breeding productivity varying from 0 – 24 chicks over two years and up to four breeding attempts. We found some evidence for relationships between telomere length and current parental effort, but no evidence that relative telomere length was related to future fecundity, survival (return rate) or cumulative (2 year) breeding productivity. Our data therefore suggest that telomere length in female starling might better reflect individual breeding and metabolic capacities than reproductive costs, even if we cannot rule out the idea that there may be adverse effects of reproduction on telomeres. However, we think that those costs actually reflect the physiological inability of LQ individuals to sustain self-maintenance and reproduction energy demands in parallel. Our data highlight the large within- and among-individual variance in telomere dynamics, as has been found in a free-ranging mammals (Fairlie, Holland, Pikington *et al.* 2016), which is difficult to reconcile with a strictly telomere-driven determination of individual quality from birth onwards. If telomeres are vulnerable to life and stress events, we need to better understand the implication of direct telomere maintenance mechanisms (*i.e.* telomerase activity, costs of maintenance) or of metabolic processes (i.e. lipid metabolism) in individual quality.

## Acknowledgements

We would like to thank David Davis and the whole Davis Family for their continued support for our European starling work on the Davistead Farm in Langley, BC. Allison Cornell provided invaluable assistance with fieldwork. We thank C. Saraux for her advice on Cq analysis.

## Authors’ Contributions

T.D.W. and M.F. collected the data and M.F. run all the physiological analyses, V.F. and S.Z. extracted the DNA and ran the qPCR measurements of telomere length, V.F., S.Z. and F.C. analysed the qPCR data, F.C. and T.D.W. did the statistical analyses, and T.D.W. and F.C. drafted the final manuscript based on a Master II report of V.F.

## Data Accessibility

On request addressed to T.D. Williams.

